# Hemagglutinin stalk-reactive antibodies interfere with influenza virus neuraminidase activity by steric hindrance

**DOI:** 10.1101/400036

**Authors:** Yao-Qing Chen, Linda Yu-Ling Lan, Min Huang, Carole Henry, Patrick C. Wilson

## Abstract

Hemagglutinin (HA) stalk-reactive antibodies are the basis of several current “one-shot” universal influenza vaccine efforts because they protect against a wide spectrum of influenza virus strains. The appreciated mechanism of protection by HA-stalk antibodies is to inhibit HA stalk reconfiguration, blocking viral fusion and entry. This study shows that HA stalk-reactive antibodies also inhibit neuraminidase (NA) enzymatic activity, prohibiting viral egress. NA inhibition (NI) is evident for an attached substrate but not for unattached small molecule cleavage of sialic acid. This suggests that the antibodies inhibit NA enzymatic activity through steric hindrance, thus limiting NA access to sialic acids when adjacent to HA on whole virions. Consistently, F(ab’)2 fragments that occupy reduced area without loss of avidity or disrupted HA/NA interactions show significantly reduced NI activity. Notably, HA stalk binding antibodies lacking NI activity were unable to neutralize viral infection via microneutralization assays. This work suggests that NI activity is an important component of HA-stalk antibody mediated protection.

**Summary:** This study reports a new mechanism of protection that is mediated by influenza hemagglutinin-stalk reactive antibodies: inhibition of neuraminidase activity by steric hindrance, blocking access of neuraminidase to sialic acids when it is abutted next to hemagglutinin on whole virions.

## Introduction

Influenza is an acute respiratory illness that causes epidemics and pandemics in the human population, causing up to 640,000 deaths annually worldwide (Iuliano et al., 2017). The influenza virus particle contains two major surface glycoproteins, hemagglutinin (HA) and neuraminidase (NA). HA is a tetramer protein that contains a globular head and a stalk domain. The head domain mediates binding to host cellular receptor sialic acids, while the stalk domain fuses the virus and host cell membranes to allow the introduction of the influenza virus genes. The NA protein is essential for cleaving terminal sialic acid residues present on host glycoproteins (Figure 1A), allowing the ready dispersal of the newly generated virus (Matrosovich et al., 2004; Palese and Compans, 1976).

**Figure 1.**
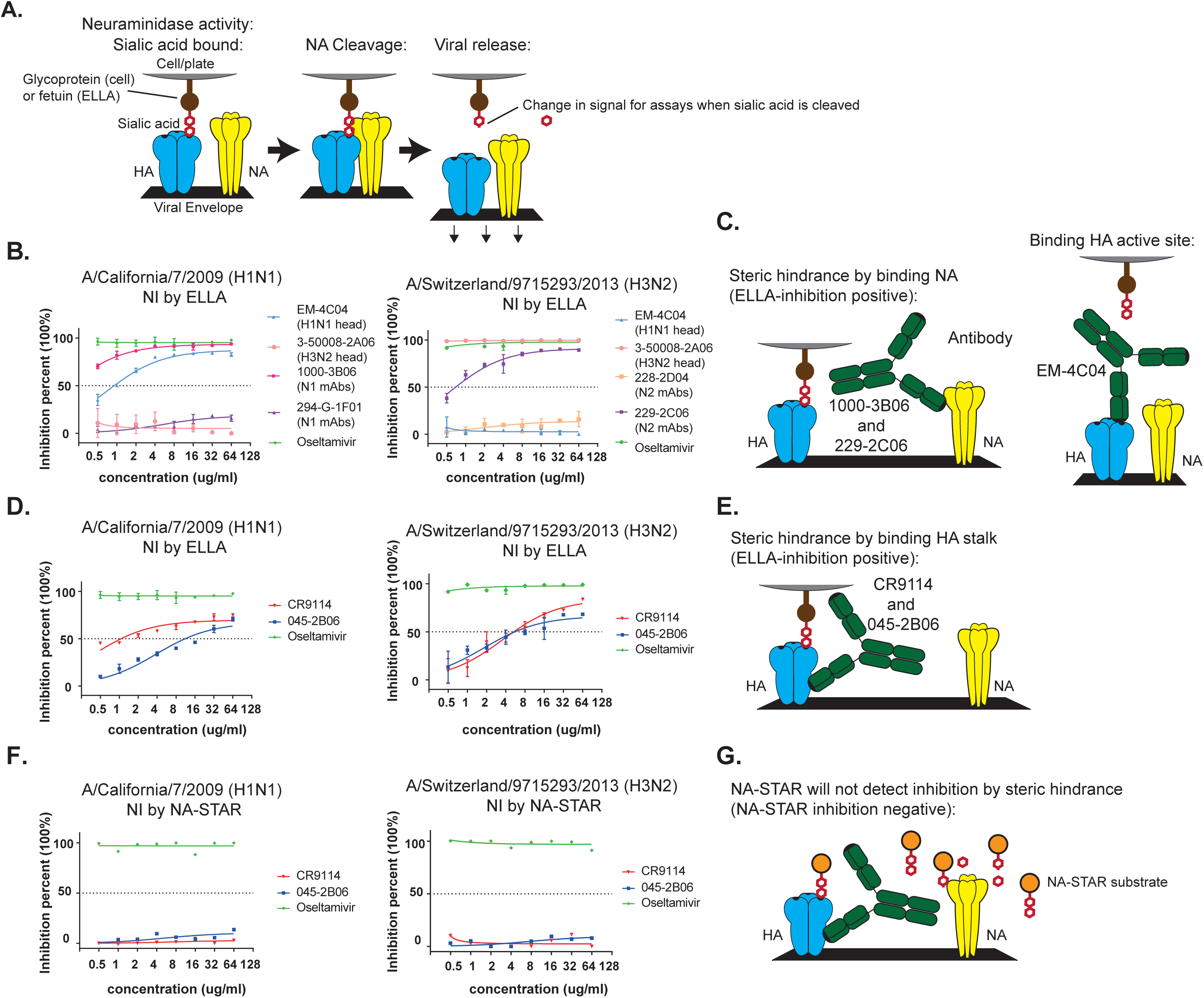
HA stalk-reactive mAbs inhibit influenza virus NA activity. **(A)** Model of NA activity to cleave bound sialic acids to release newly-made virus particles. Note that the cleaved sialic acid substrates provide the change in signal (reduced) for the ELLA and NA-STAR assay readouts. **(B)** Examples of HA head-reactive mAbs and NA-binding mAbs with NI activity are provided to illustrate inhibition of NA enzymatic activity via ELLA assays against A/California/7/2009 (H1N1) virus and A/Switzerland/9715293/2013 (H3N2) viruse. **(C)** Model for how HA head-reactive mAb and NA-reactive mAb inhibit NA from cleaving off the sialic acid from the host cell receptors. **(D)** HA stalk-reactive mAbs were tested for inhibiting NA enzymatic activity via ELLA assays against A/California/7/2009 (H1N1) virus and A/Switzerland/9715293/2013 (H3N2) virus. **(E)** Model for how HA stalk-reactive mAbs inhibit NA from cleaving off the sialic acid from the host cell receptors via steric hindrance. **(F)** HA stalk-reactive mAbs were tested for inhibiting NA enzymatic activity via NA-STAR assays against A/California/7/2009 (H1N1) virus and A/Switzerland/9715293/2013 (H3N2) virus. **(G)** Models for why NA-reactive but not HA stalk-reactive mAbs can inhibit NA from cleaving the sialic acid in the NA-STAR assays. Oseltamivir was used as a positive control. Results are shown as mean ± s.d. of three independent experiments.

Antibodies are a major means of protection from influenza virus infections. Antibody-mediated immunity is the basis of current vaccines and most efforts for improving immunity to influenza. Influenza virus vaccinations induce antibodies that predominantly target the immunodominant globular head of influenza HA that blocks viral attachment to prevent infection (Corti et al., 2010; Wrammert et al., 2011). However, immunity to the HA-head domain is highly susceptible to influenza antigenic drift, or viral mutation, introducing novel amino acids and glycosylation sites that allow the virus to evade existing immunity. The stalk is a more conserved domain, allowing antibodies that target this region to neutralize a wide spectrum of influenza virus subtypes (Corti et al., 2011; Ekiert et al., 2009; Neu et al., 2016; Okuno et al., 1993). The current appreciated mechanism of protection by HA stalk-reactive antibodies (Abs) is to lock the hemagglutinin trimer in a pre-fusion conformation, preventing pH triggered conformational changes upon viral uptake into endocytic compartments. This conformational change exposes the fusion peptide that mediates fusion of the viral membrane to the host cell membrane and subsequent introduction of the viral genome (Wu and Wilson, 2018). Because of the highly conserved epitopes within the HA-stalk domain, induction of antibodies targeting this domain are the basis of several universal influenza vaccine concepts and ongoing clinical trials. Herein, we describe an additional mechanism of protection that is mediated by HA-stalk binding antibodies: the inhibition of NA activity through steric hindrance blocking access to HA-bound sialic acid.

We used a panel of well-characterized HA stalk-reactive mAbs to explore how the mAbs interfere with NA enzymatic activity as measured by enzyme linked lectin assay (ELLA) or the NA-STAR assay. ELLA uses sialated glycoprotein fetuin that is immobilized as a substrate to measure NI. This immobilized substrate will identify antibodies that inhibit either directly by binding close to the enzymatic active site or that sterically inhibits NA from abutting close enough to HA to cleave the sialic acid (Couzens et al., 2014). Conversely, the NA-STAR assay uses a small, soluble chemiluminescent substrate, and so more explicitly distinguishes antibodies that directly inhibit the enzymatic activity of NA by binding close to the enzymatic site (Chen et al., 2018; Nguyen et al., 2010). Herein, we show that HA stalk-reactive mAbs inhibit the enzymatic activation of influenza A viruses by ELLA but not in NA-STAR assays, supporting a steric-inhibition model. Moreover, we detected reduced NI activity for the F(ab’)2 fragments from the stalk-reactive mAbs compared to the same whole antibodies, providing direct evidence that anti-HA stalk antibodies are sterically limiting NA access to sialic acids. The NI activity of HA-stalk binding but not NA-binding antibodies is disrupted by dissociating HA and NA from virions, further supporting steric-hindrance based mechanism of NI. Notably, only antibodies against the HA stalk with NI activity, including 2 with high affinity binding, were able to mediate microneutralization. These findings suggest that an important component of protection by anti-HA stalk reactive antibodies is via steric-inhibition of NI activity.

## Results

### Stalk-reactive mAbs interfere with influenza A virus NA activity

It has been shown that a subset of NA-reactive antibodies bind NA outside of the enzymatic site and inhibit sialic acid cleavage only in ELLA but not NA-STAR assays (Chen et al., 2018). Two examples of these antibodies are indicated in Figures 1B (1000-3B06 against H1N1 influenza, and 229-2C06 against H3N2 influenza). The mechanism of action of these antibodies is likely through steric inhibition of NA from accessing sialic acid molecules bound by HA (Figure 1C, left). Similarly, antibodies to epitopes on the outside periphery of the globular head of HA that do not inhibit hemagglutination, and so do not bind the HA receptor-binding domain, can nonetheless mediate protection by inhibiting viral egress similar to NI (Dreyfus et al., 2012). Thus, we hypothesized that antibodies to the stalk region of HA, which will occupy space on the side of the molecule, could similarly mediate NI activity, compounding the mechanism of protection mediated to include both inhibition of fusion and inhibition of egress. As expected (Kosik and Yewdell, 2017), antibodies binding the HA-head region of H1N1 and H3N2 influenza virus strains robustly inhibit NA activity by ELLA (Figure 1B, EM-4C04 against H1N1 influenza and 3-50008-2A06 against H3N2 influenza) because HA cannot bind the sialic acid on fetuin, and so NA is not in enough proximity for enzymatic activity (Figure 1C, right). To determine if HA-stalk antibodies elicit NI activity, we tested the well-characterized mAb 045-2B06 (Henry Dunand et al., 2015) that binds the HA-stalk regions of group 1 and 2 influenza A HA molecules and mAb CR9114 that binds the HA-stalk of both A and B influenza strain HAs (Dreyfus et al., 2012). Both of these antibodies neutralize influenza *in vitro* and protect mice *in vivo* upon lethal challenge with influenza and can competitively inhibit the binding of the other (Henry Dunand et al., 2015), demonstrating that they bind similar protective HA-stalk epitopes. We find that both 045-2B06 and CR9114 provide NI activity against both A/California/7/2009 (H1N1) and A/Switzerland/9715293/2013 (H3N2) by ELLA assay (Figure 1D). The likely mechanism for this inhibition is steric hindrance in which the antibodies block access of NA to the sialic acids mound by HA (Figure 1E). Conversely, neither 045-2B06 nor CR9114 had NI-activity to cleave the small, unattached substrate in the NA-STAR assay (Figure 1F), suggesting a mechanism of steric hindrance (compare Figures 1E versus 1G). Our results demonstrate that human antibodies reactive to the HA-stalk inhibit the enzymatic activity of NA protein on H1N1 and H3N2 influenza virus particles.

### Stalk-reactive mAbs interfere with NA activity by steric hindrance

Because the antibodies inhibited NI activity on the attached substrate of the ELLA assay but not on the free substrate of the NA-STAR assay, as depicted in Figures 1D and 1F, it appears the mechanism of inhibition is likely steric hindrance of NA access to the sialic acid when bound by HA. To investigate this possibility directly, we generated F(ab’)2 molecules from the 045-2B06 and CR9114 mAbs to reduce the size of the molecules, and thus the degree of steric hindrance, without affecting the avidity of binding. The F(ab’)2 concentration of protein were also increased to ensure equivalent binding of the F(ab’)2 molecules with that of the mAbs (Figure 2A and 2B). Comparison via the ELLA assay showed that indeed the F(ab’)2 molecules had significantly reduced NI activity in comparison to the whole mAbs against both H1N1 and H3N2 influenza viruses (Figure 2C). Thus, while not completely allowing access of NA to the sialic acid, the smaller antibody fragments allowed increased NA access to the substrate (Figure 2D), supporting a steric hindrance model of NI by HA-stalk binding antibodies. As a final verification that antibodies to the HA-stalk were inhibiting NA through steric hindrance, we dissociated the HA and NA molecules from the virions, thus allowing NA to access the sialic acid substrates completely independently of HA, regardless of the presence of antibody. For this analysis we performed the ELLA assay in the presence of 1% of Triton X-100 to disrupt the viral envelope lipid bilayer, releasing HA and NA similarly to seasonal split influenza virus vaccine production (Gross et al., 1981). The detergent treatment completely abrogated NI activity of all the HA-stalk-reactive and HA-head-reactive mAbs against both H1N1 and H3N2 virus strains, further supporting the steric hindrance model for NI activity (Figure 3A and 3B). However, the NA-reactive antibodies were able to inhibit NI activity, demonstrating integrity of the assay after detergent treatment. In total, these various results demonstrate that in addition to the well-appreciated mechanism of action disrupting viral entry, antibodies to the HA-stalk region also inhibit the access of NA to sialic acid by steric hindrance and thus reduce viral egress.

**Figure 2.**
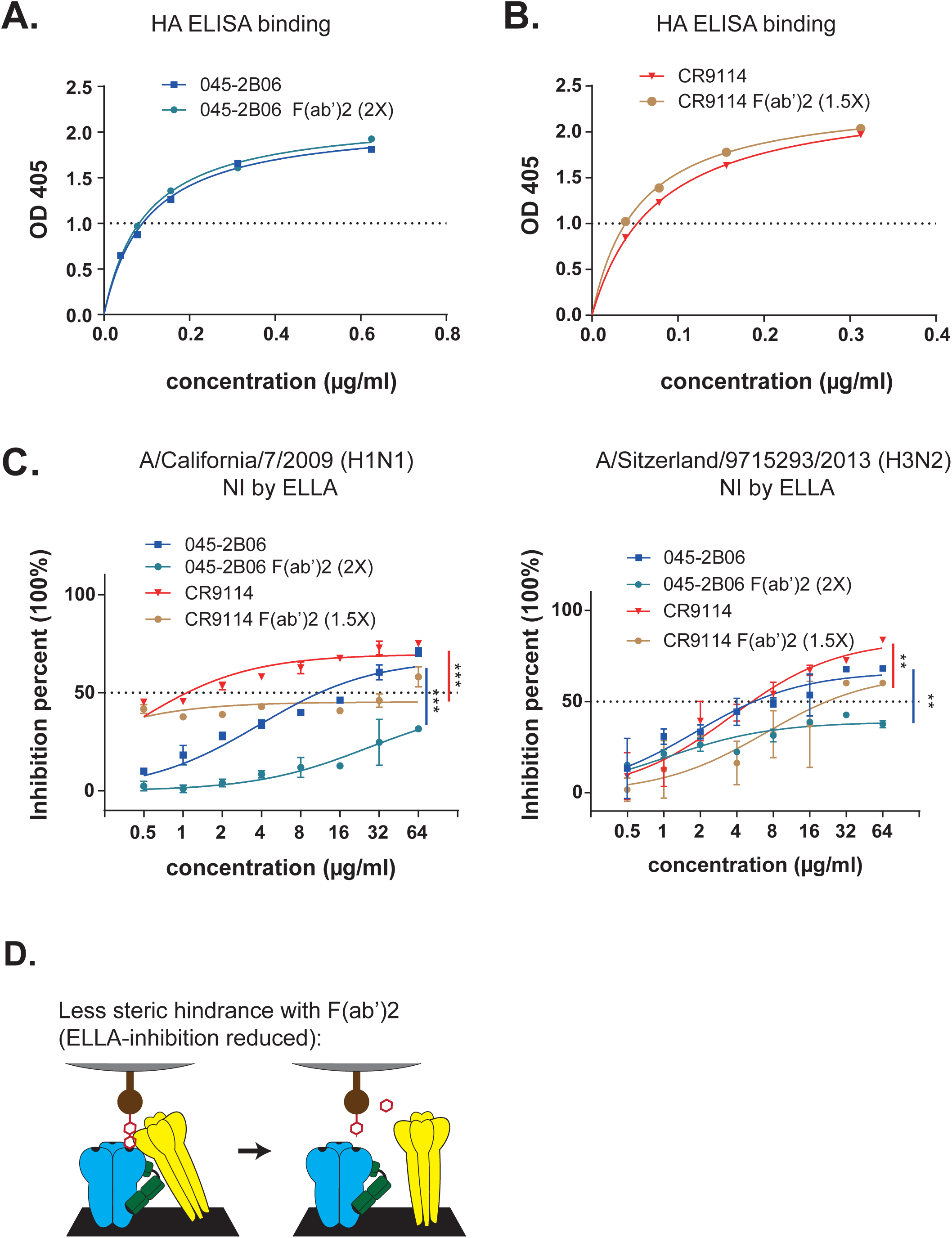
F(ab’)2 molecules from stalk-reactive mAbs have reduced interference of virus NA activity by steric hindrance. **(A)** Binding avidity of 045-2B06 and its F(ab’)2 fragment against A/California/7/2009 rHA. **(B)** Binding avidity of CR9114 and its F(ab’)2 fragments against A/California/7/2009 rHA. **(C)** HA stalk-reactive mAbs and their F(ab’)2 fragments were tested for inhibiting NA enzymatic activity via ELLA assays against A/California/7/2009 (H1N1) virus and A/Switzerland/9715293/2013 (H3N2) virus. **(D)** Model for why F(ab’)2 fragments of HA-stalk reactive mAbs have reduced inhibition of NI in ELLA assays. Results are shown as mean ± s.d. of three independent experiments.

**Figure 3.**
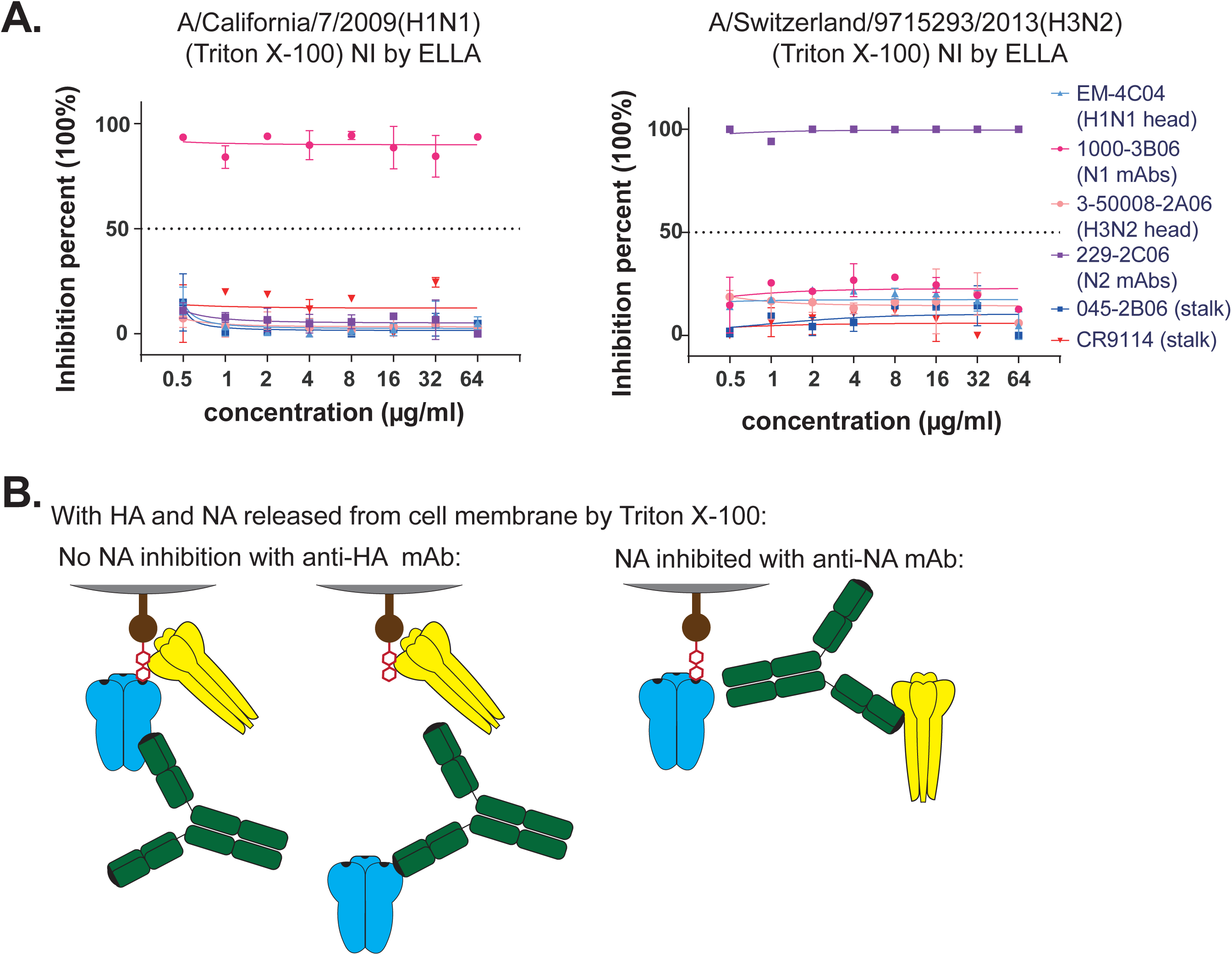
HA-stalk reactive mAbs cannot interfere with influenza virus NA activity if HA and NA are dissociated from the viral envelope. A/California/7/2009 (H1N1) virus and A/Switzerland/9715293/2013 (H3N2) virus were treated with 1% TritonX-100 for 1h at 37C **(A)** HA-stalk and -head reactive mAbs as well as NA-reactive mAbs were tested for NI activity via ELLA assays against TritonX-100 treated A/California/7/2009 (H1N1) virus and A/Switzerland/9715293/2013 (H3N2) virus. **(B)** Model for why HA-reactive mAbs cannot disrupt NA activity whereas NA-reactive mAbs can when the HA and NA glycoproteins are dissociated from the viral envelope. Results are shown as mean ± s.d. of two independent experiments.

### Stalk-reactive mAbs have varying capacities for NA enzymatic inhibition activity that appears important for potency

Various HA-stalk reactive antibodies have been shown to have distinct amino acids involved in binding, various angles of interaction, and differences in specificity as indicated by the spectrum of influenza strains bound (Wu and Wilson, 2018). Thus it is predicted that not all HA-stalk binding antibodies will elicit NI activity to the same degree. Therefore we tested a panel of 13 human mAbs that were verified to bind HA-stalk epitopes (Andrews et al., 2015) for NI activity by ELLA. Criteria for being classified as HA-stalk reactive included competitive binding with structure-verified HA-stalk reactive antibodies including CR9114 (Dreyfus et al., 2012) and SC70-F02 (Nachbagauer et al., 2018), as well as binding in a pH dependent fashion as the HA-stalk epitope is disrupted at low pH (Andrews et al., 2015). The majority, but not all, of the anti-HA stalk antibodies tested could inhibit NA activity, including: 69% (9 of 13) that are reactive with an H1N1 influenza virus (Figure 4A and 4C) and 3 of 4 on hand that bind an H3N2 influenza virus (Figure 4B and 4D). Notably, none of these antibodies had NI activity using the NA-STAR assay (Figure 4E-H), suggesting that as with CR9114 and 045-2B06, they all inhibit NI via steric interference of NA to sialic acid. Importantly, after testing the panel of mAbs by microneutralization (MN) assay, we noted that only those antibodies that have NI activity were able neutralize viral infection *in vitro* (Figure 5A and 5B). While two of the antibodies that lack MN capacity were low avidity, possibly explaining the lack of both MN and NI activities, the other two were of equal avidity to the set of HA stalk binding antibodies that did inhibit NI and neutralize (Figure 5C). The lack of NI and MN activity of two of the HA-stalk binding antibodies that have high avidity suggests that the epitope is bound in an atypical fashion. Thus we considered if these antibodies were of the stereo-typical variety with restricted variable gene repertoires, biased for VH1-69 and VH1-18 immunoglobulin heavy chain gene usage. Notably, while most of the NI+ antibodies appear to be stereotypical (Figure 6A), none of the NI-negative anti-HA stalk antibodies used the most typical anti-HA stalk VH gene, VH1-69 (Figure 6B). Also, one of the high avidity but NI-negative antibodies used VH3-23, which is indeed atypical, while the other high avidity NI-negative antibody was encoded by VH1-18, which is a common stalk-related gene that is highly similar to VH1-69. There were no other features of the immunoglobulin gene repertoire that were distinct between NI-positive and NI-negative antibodies (Figure 6C-6F). These studies demonstrate that while the majority of HA-stalk reactive antibodies inhibit NA activity a subset do not and the capacity for NI appears to be important for potency.

**Figure 4.**
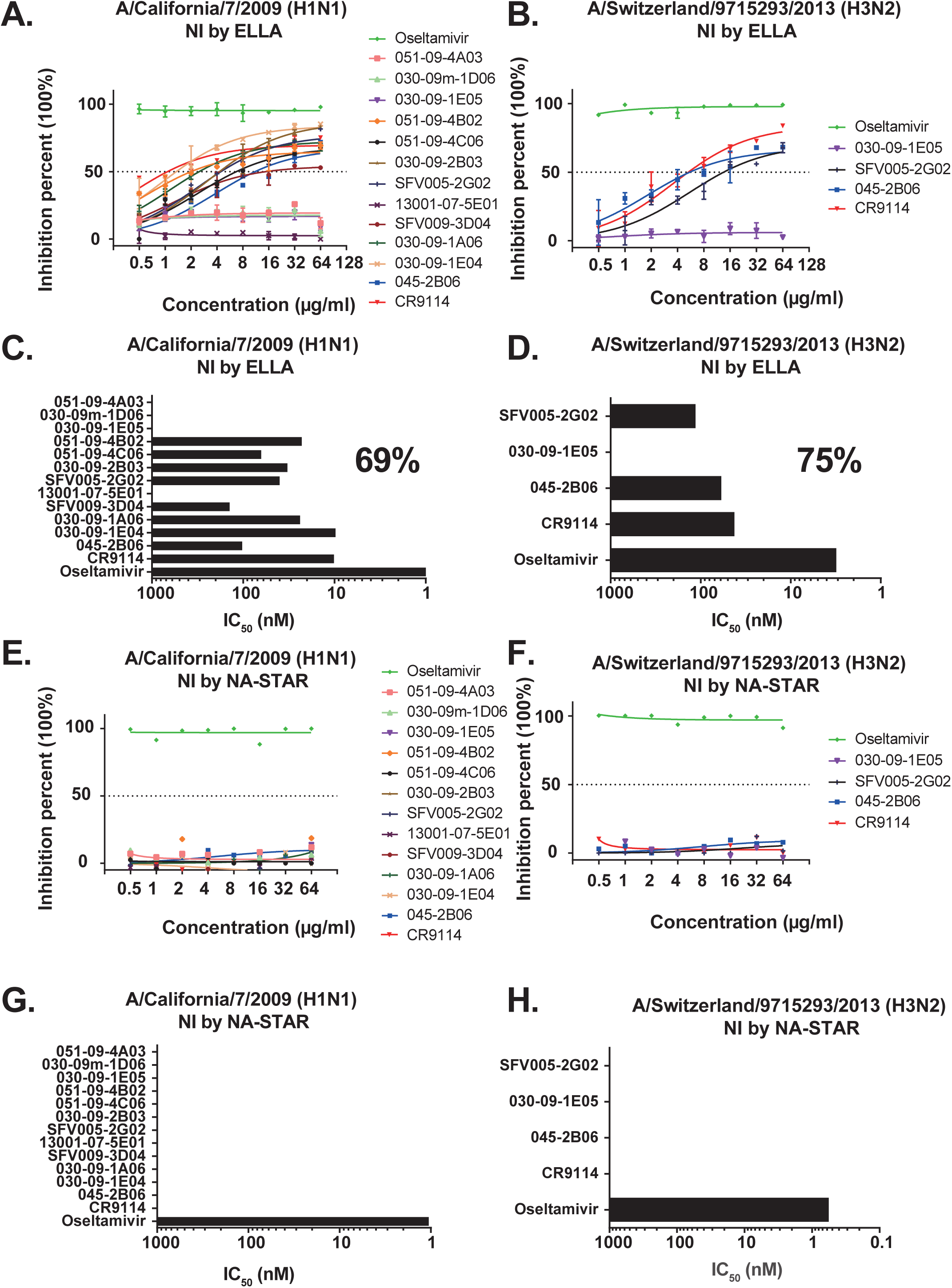
Stalk-reactive mAbs affect virus NA activity differently (A and C) A set of HA stalk-reactive mAbs were tested for NI activity via ELLA assays against A/California/7/2009 (H1N1) virus. Oseltamivir was used as a positive control. **(B and D)** A set of HA stalk-reactive mAbs were tested for NI activity via ELLA assays against A/Switzerland/9715293/2013 (H3N2) virus. are represented as half-maximum inhibitory concentration IC_50_ (nM). **(E and G)** A set of HA stalk-reactive mAbs were tested for NI activity via NA-STAR assays against A/California/7/2009 (H1N1) virus. **(F and H)** A set of HA stalk-reactive mAbs were tested for NI activity via NA-STAR assays against A/Switzerland/9715293/2013 (H3N2) virus. Data in panels C, D, G, and H represent half-maximum inhibitory concentrations or IC_50_ (nM). Oseltamivir was used as a positive control for all assays. Data are representative of two independent experiments.

**Figure 5.**
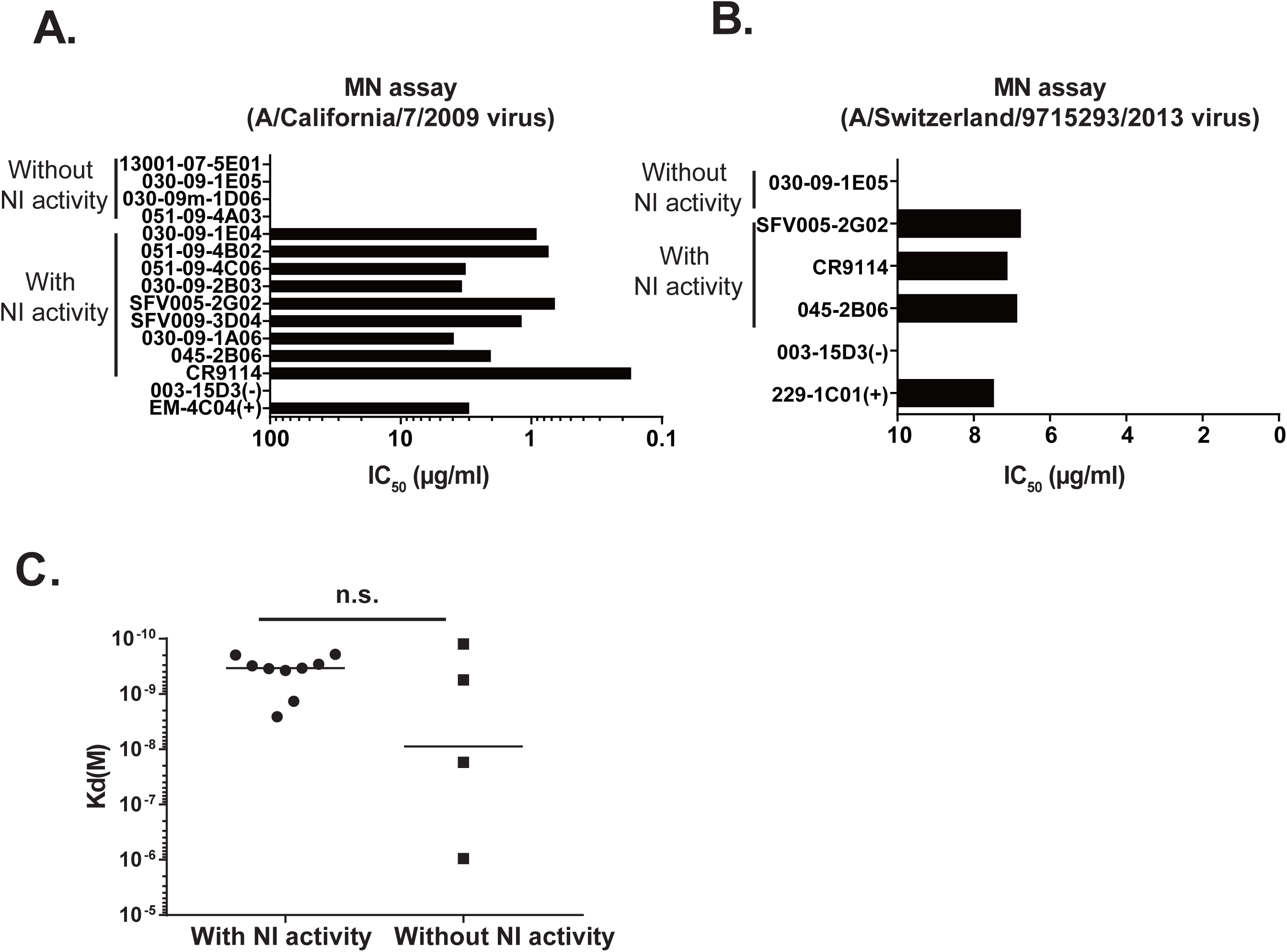
NI activity of influenza HA-stalk reactive antibodies correlates with microneutralization capacity. **(A and B)** HA stalk-reactive mAbs with or without NI activity were tested for neutralization in vitro microneutralization (MN) assays using (A) A/California/7/2009 (H1N1) virus and (B) A/Switzerland/9715293/2013 (H3N2) virus. Data are represented as half-maximum inhibitory concentrations or IC_50_ (µg /ml). Positive control HA-reactive mAbs EM-4C04 (anti-H1N1) and 229-1C01 (anti-H3N2) bind HA and neutralize these influenza virus strains. Influenza-non-reactive mAb 003-15D3 was used as a negative control in the assays. **(C)** HA stalk-reactive mAbs were tested for binding affinity against HA recombinant protein. Binding avidities (K_D_) were estimated by Scatchard plot analyses of ELISA data. Bars indicate median values. Data are representative of two independent experiments. n.s., not significant.

**Figure 6.**
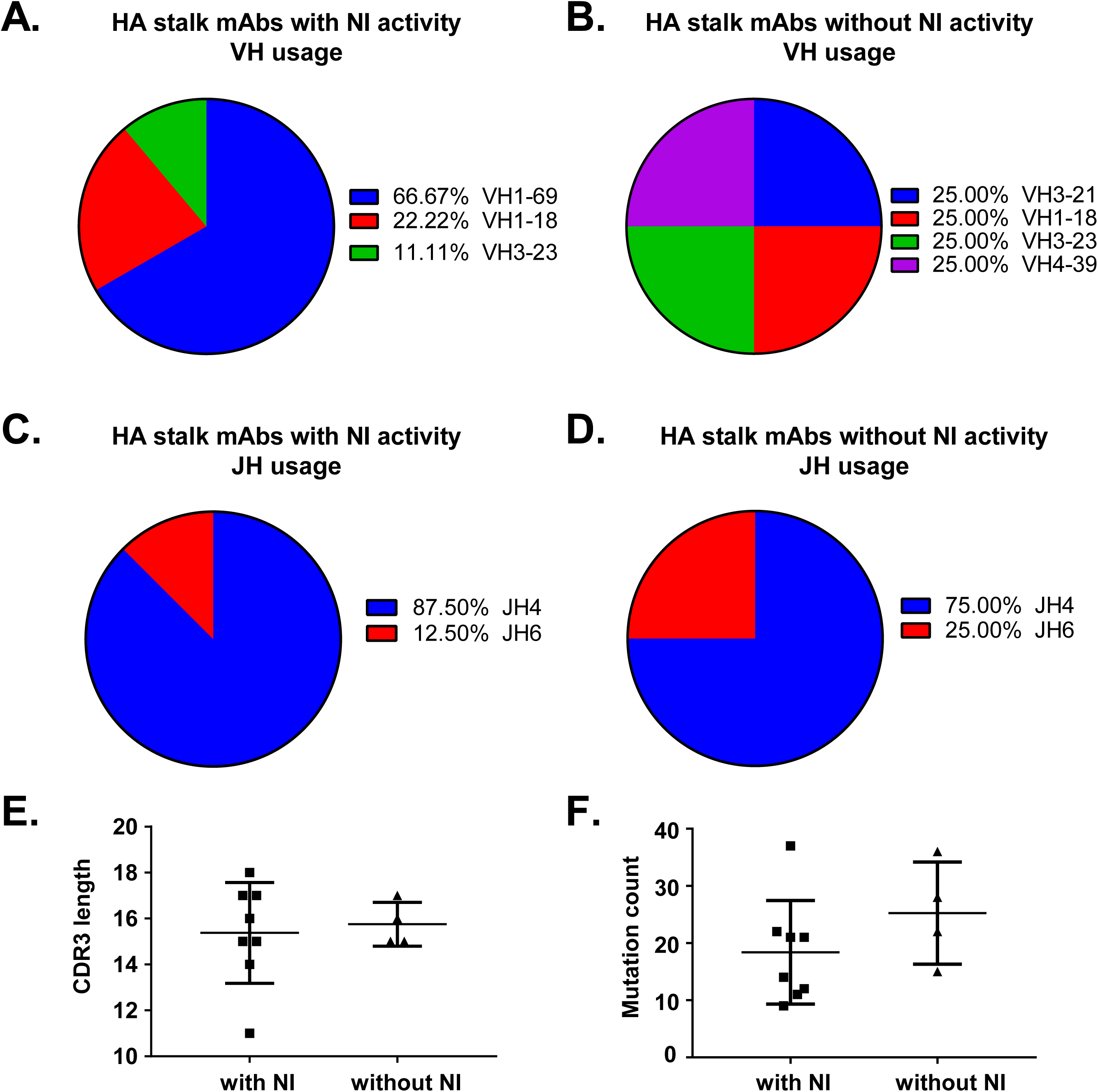
Heavy chain gene features of HA stalk-reactive mAbs with or without NI activity. **(A and B)** Usage of VH immunoglobulin genes by (A) HA stalk-reactive B cells with NI activity and (B) HA stalk-reactive B cells without NI activity. **(C and D)** Usage of JH immunoglobulin genes by (C) HA stalk-reactive B cells with NI activity and (D) HA stalk-reactive B cells without NI activity. **(E)** CDR3 length of HA stalk-reactive mAbs with and without NI activity, data are represented as mean ± SD. (F) Total heavy chain amino acid mutations for HA stalk-reactive mAbs with and without NI activity based on analysis using the NCBI IgBlast tool (https://www.ncbi.nlm.nih.gov/igblast/), data are represented as mean ± SD.

## Discussion

The development of a “one-shot” universal influenza vaccine that will provide long-term or improved immunity to influenza epidemics and pandemics is currently a major focus of the biomedical community (Paules et al., 2017a; Paules et al., 2017b). The HA stalk is an important target for the ongoing design of universal influenza vaccines and for recent and ongoing clinical trials because the protective epitopes in this portion of HA are highly conserved across many influenza subtypes (Henry et al., 2018; Krammer et al., 2018; Wu and Wilson, 2018). While the appreciated mechanism of action of HA-stalk binding antibodies is to inhibit viral entry via disrupting the HA conformational change required for viral-envelope to host cell membrane fusion, there was one recent report using a microscopy-based assay suggesting the anti-HA stalk antibodies also block influenza virus particle release (Yamayoshi et al., 2017). Herein, we show that antibodies to the HA-stalk region inhibit the activity of NA through steric hindrance on whole virions, adding to the mechanisms of protection mediated by this broadly-reactive class of antibodies, and providing a mechanism for the previously observed inhibition of viral release.

It is notable that not all antibodies binding the HA-stalk region can inhibit NI activity, although for the collection of anti-stalk antibodies tested for this study, NI activity correlated perfectly with MN activity. Thus, NI activity is in the least an important correlate of neutralization-capacity, or at most a critical component of the activity of anti-HA stalk binding antibodies. In future studies, it will be important to distinguish the relative contributions of inhibiting viral entry versus viral egress for HA-stalk reactive antibodies in protection.

A second interesting observation that should be evaluated is that there is a potential correlation between the capacity for NI activity and the type of HA-stalk antibody elicited in that none of the NI-negative antibodies binding the HA stalk were encoded by the stereotypical VH1-69 gene. A more substantial evaluation of many anti-stalk antibodies for various properties and structural studies to evaluate the particular contact residues and angle of binding for stalk-binding antibodies that are NI will provide insight into this issue. Identifying stereotypical features for anti-HA stalk antibodies with NI activity could inform on universal influenza vaccine design by, for example, providing an impetus to develop germline boosting prime-boost strategies to elicit a particular class of anti-HA stalk antibodies preferentially.

In conclusion, this study identifies NI inhibition as a new and additional mechanism of action for the important class of anti-influenza antibodies that bind highly-conserved epitopes on the HA stalk.

## Materials and Methods

### Cells and viruses

Both the Human Embryonic Kidney (HEK) 293T cells and Madin-Darby Canine Kidney (MDCK) cells were obtained from ATCC. The 293T cells were maintained at 37°C with 5% CO2 in Advanced DMEM cell medium with 2% Ultra-Low IgG FBS (Gibco), 2 mM GlutaMAX (Gibco), plus penicillin and streptomycin (100 mg/ml; Gibco). The MDCK cells were maintained at 37°C with 5% CO2 in DMEM cell medium with 10% FBS (Gibco), 1% L-Glutamine (Gibco) and 1% Antibiotic-Antimycotic (Gibco) with All influenza virus stocks (A/California/7/2009 H1N1 and A/Switzerland/9715293/2013 H3N2) used for the assays were freshly grown in specific pathogen free (SPF) eggs, harvested, purified, titered and stored at −80°C.

### Recombinant monoclonal antibody expression and purification

Antibodies were generated as previously described (Andrews et al., 2015; Henry Dunand et al., 2016; Smith et al., 2009; Wrammert et al., 2008). Briefly, the VH, Vk or Vλ genes amplified from each single cell were cloned into IgG1, Igk or Igλ expression vectors as previously mentioned. 9 µg of each paired heavy and light chain plasmid DNA were transfected into the 293 cells using PEI (name of the brand) and incubated overnight. The next day, the transfection media was aspirated from each plate and replaced by 25 ml of PFHM-II (Gibco). Secreted mAbs were purified from the supernatant using protein A beads 4-5 days later. The mAbs was further concentrated and buffer exchanged with PBS (pH 7.4) using Amicon Ultra centrifugal filter units (30 kDa cutoff; Millipore). The final protein concentration was determined using a NanoDrop device (Thermo Scientific).

### Preparation of F(ab’)2 monoclonal antibody fragments

The whole antibodies CR9114 and 045-2B06 were reduced to a F(ab’)2 fragment using the Pierce™ F(ab’)2 digestion kit (Cat#: 44988, Thermo Fisher) according to the manufactures instructions.

### Enzyme linked immunosorbent assay (ELISA)

High-protein binding microtiter plates (Costar) were coated with recombinant HAs or NAs at 1µg/ml in phosphate buffered saline (PBS) overnight at 4°C. After blocking, serially diluted (3 fold) antibodies starting at 10µg/ml were incubated for 1 h at 37°C. Horse radish peroxidase (HRP)-conjugated goat anti-human IgG antibody diluted 1:1000 (Jackson Immuno Research) was used to detect binding of mAbs, and was developed with Super Aquablue ELISA substrate (eBiosciences). Absorbance was measured at 405nm on a microplate spectrophotometer (BioRad). To standardize the assays, antibodies with known binding characteristics were included on each plate and the plates were developed when the absorbance of the control reached 3.0 OD units (Denise Lau et al., 2017). To determine the binding of F(ab’)2 fragment by ELISA, an HRP-conjugated goat anti-kappa antibody (SouthernBiotech) was used as a secondary antibody.

### NA enzyme linked lectin assay (ELLA)

ELLAs were performed as previously described (Westgeest et al., 2015). Flat-bottom 96-well plates (Thermo Scientific) were coated with 100 μl of fetuin (Sigma) at 25 µg/ml overnight at 4°C. Antibodies were serially diluted (twofold) in Dulbecco’s phosphate-buffered saline (DPBS) with 0.05% Tween 20 and 1% BSA (DPBSTBSA), then incubated in duplicate fetuin-coated plates with an equal volume of the selected virus particles dilution in DPBSTBSA. These plates were incubated for 18 h at 37°C and washed six times with PBS with 0.05% Tween 20, and 100 μl/well of HRP-conjugated peanut agglutinin lectin (PNA-HRPO, Sigma–Aldrich) in DPBSTBSA was added for 2h at RT in the dark. The plates were washed six times and were developed with Super Aquablue ELISA substrate (eBiosciences). Absorbance was read at 405nm on a microplate spectrophotometer (BioRad). Data points were analyzed using Prism software and the 50% inhibition concentration (IC_50_) was defined as concentration at which 50% of the NA activity was inhibited compared to the negative control.

### NA-STAR assay

The NA-STAR assay was performed according to the Resistance Detection Kit manufacturer’s instructions (Applied Biosystems, Darmstadt, Germany) (Nguyen et al., 2010). In brief, 25 μl test mAbs in serial twofold dilutions in NA-STAR assay buffer (26 mM 2-(N-morpholino) ethanesulfonic acid; 4 mM calcium chloride; pH 6.0) were mixed with 25 μl of 4 X IC_50_ of virus and incubated at 37°C for 30 min. After adding 10 μl of 1000× diluted NA-STAR substrate, the plates were incubated at room temperature for another 30 minutes. The reaction was stopped by adding 60 μl of NA STAR accelerator. The chemiluminescent was determined by using the DTX 880 plate reader (Beckman Coulter). Data points were analyzed using Prism software and the 50% inhibition concentration (IC_50_) was defined as concentration at which 50% of the NA activity was inhibited compared to the negative control. The final concentration of antibody (IC_50_) was determined using Prism software (GraphPad).

### NI assay in the presence of detergent

To perform NI assay in the presence of detergent, all steps remained identical to those listed above, except the following: firstly, a final concentration of 1% of Triton X-100 (Fisher Bioreagents) was added directly to the virus particles, shacked gently at 37 °C for 1 hour. During this time, antibodies were diluted using PBS with 1% Triton X-100. Prior to incubation with mAbs, the virus preparation was diluted in PBS containing 1% BSA and 1% Triton X-100. The NI assay was then performed as detailed above.

### Microneutralization assay (MN)

MN assay for antibody characterization was carried out as previously described (Henry Dunand et al., 2015). Briefly, MDCK cells were maintained in Dulbecco’s modified Eagle’s medium (DMEM) supplemented with 10% fetal bovine serum (FBS) at 37°C with 5% CO2. On the day before the experiment, confluent MDCK cells in a 96-well format were washed twice with PBS and incubated in minimal essential medium (MEM) supplemented with 1 μg/ml trypsin-TPCK. Serial 2-fold dilutions (starting concentration 64 µg/ml) of mAb were mixed with an equal volume of 100 50% tissue culture infectious doses (TCID50) virus and incubated for 1 h at 37°C. The mixture was removed and cells were cultured for 20 h at 37°C with 1X MEM supplemented with 1 μg/ml trypsin-TPCK and appropriate mAb concentration. Cells were washed twice with PBS, fixed with 80% acetone at −20°C for 1 h or overnight, washed 3 times with PBST, blocked for 30 min with 10% FBS and then treated for 30 min with 2% H2O2 at room temperature (RT). An anti-NP-biotinylated antibody (1:3000) in 3% BSA-PBS was incubated for 1h at RT. The plates were developed with Super Aquablue ELISA substrate and read at 405 nm. The signal from uninfected wells was averaged to represent 100% inhibition. Virus infected wells without mAb were averaged to represent 0% inhibition. Duplication wells were used to calculate the mean and SD of neutralization, and inhibitory concentration 50 (IC_50_) was determined by a sigmoidal dose response curve. The inhibition ratio (%) was calculated as below: (OD (Pos. Control) –OD (Sample))/ (OD (Pos. Control) –OD (Neg. Control)) × 100%. The final concentration of antibody that reduced infection to 50% (IC50) was determined using Prism software (GraphPad).

### Data analysis and statistics

GraphPad Prism 7 was used to perform all statistical analyses. Statistical significance is indicated as follows: n.s. (not significant), P > 0.05; *, P ≤ 0.05; **, P ≤ 0.01; ***. Data was presented as mean + standard deviation (SD) of triplicates.

## Author contributions

Y.-Q.C. designed and performed experiments, analyzed data, and wrote the manuscript. L.Y.-L.L., M.H. and C.H. performed experiments and revised the manuscript. P.C.W. designed and directed the project, analyzed data and wrote the manuscript.

## Acknowledgments

We thank Nai-Ying Zheng for purifying influenza viruses and experimental assistance. This project was funded in part from the National Institute of Allergy and Infectious Disease; the National Institutes of Health grant numbers U19AI082724 (P.C.W.), 5U19AI109946 (P.C.W.), U19AI057266 (P.C.W.), and the NIAID Centers of Excellence for Influenza Research and Surveillance (CEIRS) grant numbers HHSN272201400005C (P.C.W., D.J.T., and J.T.).

The authors declare no financial or commercial conflicts of interest.

